# A Rigorous Interlaboratory Examination of the Need to Confirm NGS-Detected Variants with an Orthogonal Method in Clinical Genetic Testing

**DOI:** 10.1101/335950

**Authors:** Stephen E. Lincoln, Rebecca Truty, Chiao-Feng Lin, Justin M. Zook, Joshua Paul, Vincent H. Ramey, Marc Salit, Heidi L. Rehm, Robert L. Nussbaum, Matthew S. Lebo

**Affiliations:** Invitae, San Francisco, CA; Partners HealthCare Laboratory for Molecular Medicine, Cambridge, MA; Brigham and Women’s Hospital, Boston, MA; National Institute of Standards and Technology, Gaithersburg, MD; Joint Initiative for Metrology in Biology, Stanford, CA; Center for Genomic Medicine, Massachusetts General Hospital, Boston, MA; Harvard Medical School, Boston, MA; The Broad Institute of MIT and Harvard, Cambridge, MA; Department of Medicine, University of California San Francisco, San Francisco, CA

## Abstract

Orthogonal confirmation of NGS-detected germline variants has been standard practice, although published studies have suggested that confirmation of the highest quality calls may not always be necessary. The key question is how laboratories can establish criteria that consistently identify those NGS calls that require confirmation. Most prior studies addressing this question have limitations: These studies are generally small, omit statistical justification, and explore limited aspects of the underlying data. The rigorous definition of criteria that separate high-accuracy NGS calls from those that may or may not be true remains a critical issue.

We analyzed five reference samples and over 80,000 patient specimens from two laboratories. We examined quality metrics for approximately 200,000 NGS calls with orthogonal data, including 1662 false positives. A classification algorithm used these data to identify a battery of criteria that flag 100% of false positives as requiring confirmation (CI lower bound: 98.5–99.8% depending on variant type) while minimizing the number of flagged true positives. These criteria identify false positives that the previously published criteria miss. Sampling analysis showed that smaller datasets resulted in less effective criteria.

Our methodology for determining test and laboratory-specific criteria can be generalized into a practical approach that can be used by many laboratories to help reduce the cost and time burden of confirmation without impacting clinical accuracy.

## Introduction

The use of orthogonal assays (e.g., Sanger sequencing) to confirm variants identified with next-generation sequencing (NGS) is standard practice in many laboratories to reduce the risk of delivering false positive (FP) results. Clinical NGS tests can inform significant medical decisions [1,2], and therefore confirmation is recommended by medical practice guidelines [3,4], although the details are generally left up to the laboratory [3,5]. Because clinical NGS methods often emphasize sensitivity (to avoid missing clinically important variants), FP rates can be elevated compared with those in research NGS [6]. Moreover, pathogenic variants are often technically challenging (e.g., many are located within repetitive or complex regions), which can further increase FP rates [7–9]. Confirmation assays have a monetary cost, however, and also increase the time needed to deliver results, a critical factor in many clinical situations.

Published studies examining this issue have concluded that confirmation of the highest quality NGS calls may not always be necessary [10–14]. Some of these studies [10,12] propose specific criteria for separating high-confidence true-positive (TP) variant calls from those that are possible FPs. These criteria differ from those used in filtering, the separate process of removing calls confidently believed to be false or unsupportable. The remaining intermediate-confidence calls are those that benefit from confirmatory assays, additional data review, or both to determine which are TPs and which are FPs. Unfortunately, these prior studies are generally small, in some cases proposing criteria using only one or five example FPs (Table 1). The presence of few FPs may seem reassuring but leads to significant limitations in these studies. First, because their statistical power to characterize the FP population is limited, these studies do not address the question of whether future FPs are likely to resemble the few observed in the study. Quite possibly, additional FPs could be different and thus missed by the proposed criteria. Second, most of these studies use identical datasets for training and evaluating the proposed criteria, likely making the results subject to overfitting [15]. Third, few of these studies provide statistical justification. Finally, all of these prior studies use data from individual laboratories and do not examine whether the methodologies can be generalized.

**Table 1.**
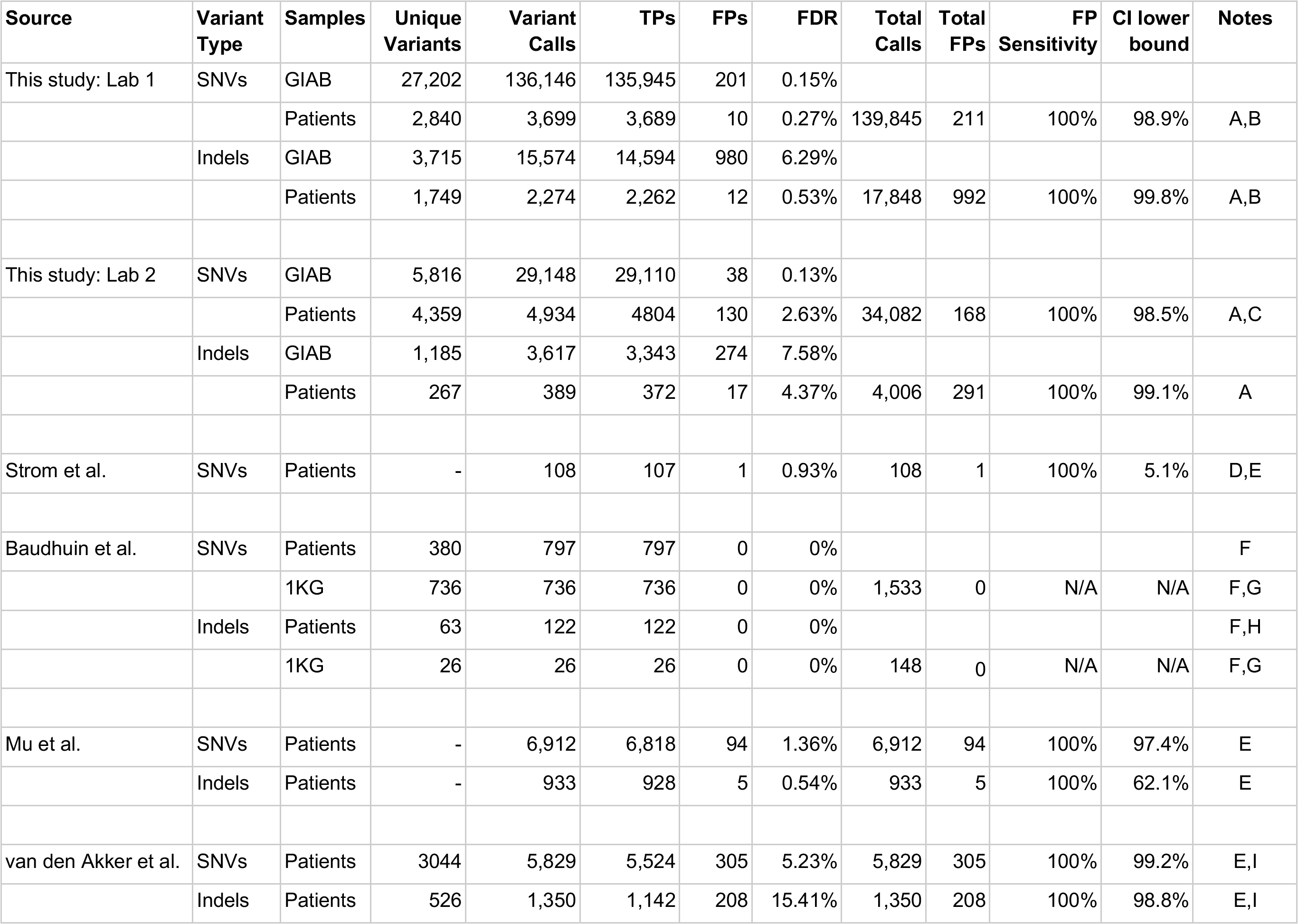

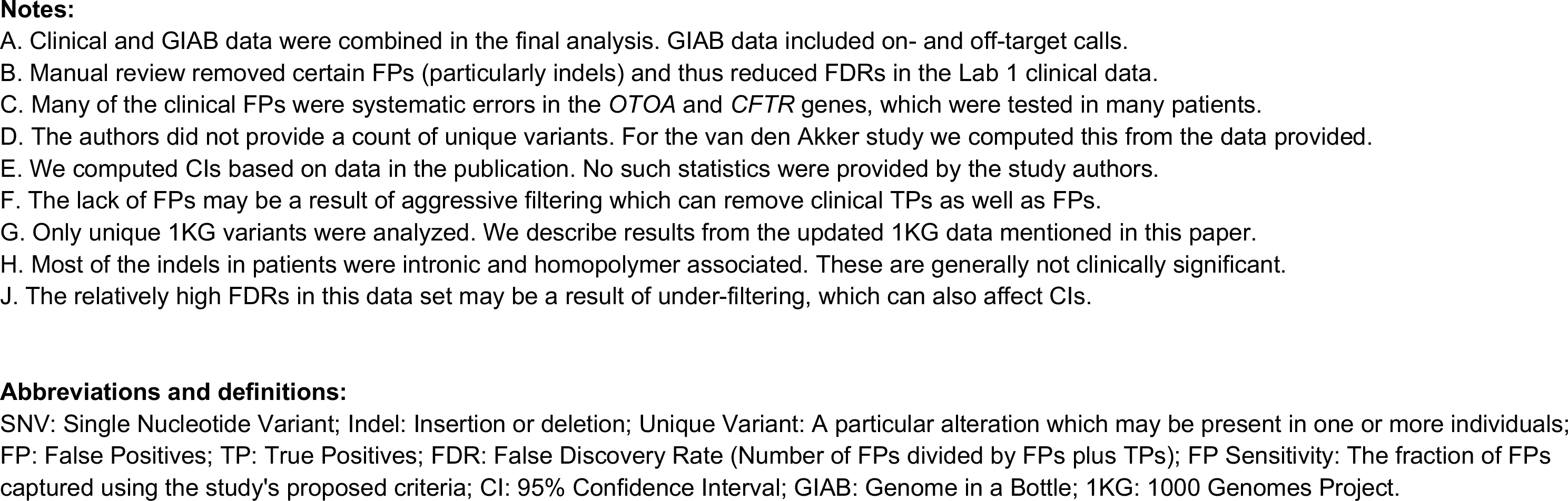

We set out to examine the role of confirmation using a set of variant calls much larger than those published previously. We combined our own sequences of five reference samples characterized by the Genome in a Bottle Consortium (GIAB) [16–18] with confirmatory data from over 80,000 clinical tests. Our methodology was applied in two clinical laboratories that use similar but not identical NGS methods. Similar to prior studies, we were able to to identify high-confidence NGS calls that do not benefit from confirmation. However, we found that a battery of criteria was necessary to capture all FPs, in contrast with the one or two metrics used by most of the prior studies. Indeed, we found that the specific criteria proposed by prior studies would miss FPs in our much larger dataset. We also found that observations of a variant as a TP can say little about its chance of being an FP in a different sample or NGS run, which indicates that prior confirmations can be an ineffective quality metric. Approaches such as ours can be used by any clinical laboratory to provide efficient, effective, and statistically justified criteria for prioritizing variant calls for confirmation.

## Methods

We compiled eight component datasets from two laboratories (Table 1) following the process illustrated in Figure 1. Key aspects of our methodology are summarized in Table 2 and are detailed here and in our Discussion below. In addition to results obtained through clinical testing, five GIAB DNA specimens were sequenced: NA12878, NA24385, NA24143, NA24149, and NA24631 (Coriell Institute, Camden, NJ). Replicates of the GIAB samples were included. NGS was performed on both the GIAB and clinical specimens by each laboratory using Illumina (San Diego, CA) 2 × 150 base pair (bp) reads as described previously [8,19,20]. Seven (Lab 1) and three (Lab 2) custom hybridization-based assays were used, each targeting 100–1000 genes. Clinically reportable target regions included, with some exceptions, protein-coding exons plus immediate flanking sequences (10–20 bp on each side). Average coverage across targets was 300-1000x or more depending on the sample, assay, and laboratory. All data used in this study passed stringent sample and run-level quality controls.

**Figure 1:**
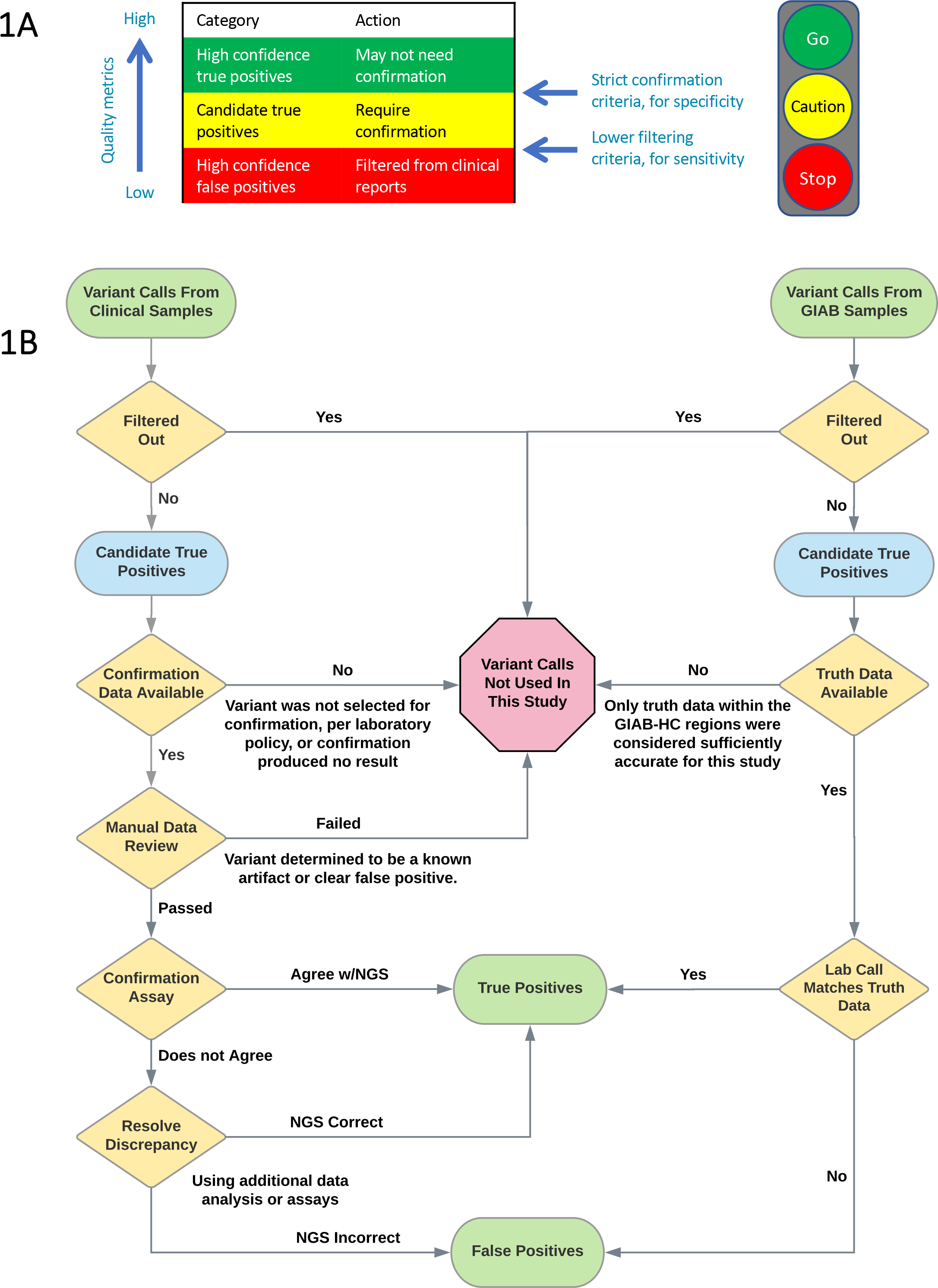
Study methodology. (A) Variant calls can be separated into high-confidence true positive (green) and intermediate-confidence (yellow) categories using strict thresholds intended to maximize the specificity of the high-confidence set. The intermediate set will contain a mixture of true and false positive calls. Our study objective was to rigorously determine test-specific criteria that separate these two categories. Variant calls that are confidently false positives (red) are typically filtered out using different criteria that emphasize sensitivity. The analogy to a traffic light is illustrative. (B) Process for collecting true positive and false positive variant calls for both the clinical and Genome in a Bottle (GIAB) specimens. Each laboratory’s clinically validated NGS assays, bioinformatics pipeline, and filtering criteria were used for both specimen types. Singlenucleotide variants and indels were collected as shown. Copy number and structural variants were excluded from this study, as were any variants with an unknown confirmation status. Filtering and manual review processes were designed to remove clearly false variant calls but not those considered even potentially true. Manual review was used only with the Lab 1 clinical data.

**Table 2.**

**Main Table 2**
| **Aspect of our methodology** | **Rationale (see text for details)** |
| --- | --- |
| Large datasets used for both SNVs and Indels | Provides confidence in the resulting criteria and helps minimize overfitting. Appropriate sizes determined by CI calculations (below). |
| Both clinical and reference (GIAB) samples employed | Greatly increases the size and diversity of our datasets, particularly in FPs. |
| Same quality filtering thresholds as used in clinical practice | Confirmation criteria can depend on filtering criteria. Using lower filters would add many FPs to our study but could result in the selection of ineffective or biased confirmation criteria. |
| Separate filtering and confirmation thresholds used | Allows high sensitivity (by keeping variants of marginal quality) and high specificity (by subjecting these variants to confirmation). |
| On- and off-target variant calls analyzed in GIAB samples | Further increases the dataset size and diversity. Off-target calls were subject to the same quality filters as on-target calls. |
| Indels and SNVs analyzed separately | Indels and SNVs can have different quality determinants. We required an adequate population of each to achieve statistical significance. |
| Partial matches considered FPs | Zygosity errors and incorrect diploid genotypes do occur and can be as clinically important as “pure” FPs. |
| Algorithm selects criteria primarily by their ability flag FPs | Other algorithms equally value classification of TPs, which may result in biased criteria, particularly as TPs greatly outnumber FPs. |
| Multiple quality metrics used to flag possible FPs | Various call-specific metrics (e.g. quality scores, read depth, allele balance, strand bias) and genomic annotations (e.g. repeats, segmental duplications) proved critical. |
| Key metric: Fraction of FPs flagged (FP sensitivity) | Other metrics, including test PPV and overall classifier accuracy can be uninformative or misleading for evaluating confirmation criteria. |
| Requirement of 100% FP sensitivity on training data | Clinically appropriate. Resulting criteria will be effective on any subset of the training data (clinical or GIAB, on-target or off-target, etc.) |
| Statistical significance metric: CI on FP sensitivity | Rigorously indicates validity of the resulting criteria: e.g.. Flagging 100% of 125 FPs demonstrates $\geq 98\%$ FP sensitivity at $p=0.05$.<br>Smaller datasets (e.g. 50 FPs) resulted in ineffective criteria. |
| Separate training and test sets employed (cross-validation) | In conjunction with large datasets, cross-validation is a crucial step to avoid overfitting which can otherwise result in ineffective criteria. |
| Prior confirmation of a variant was not used as a quality metric | Successful confirmations of a particular variant can indicate little about whether future calls of that same variant are true or false. |
| All variants outside of GIAB-HC regions require confirmation | Outside of these regions we have too little confirmatory data to prove whether our criteria are effective or not. |
| Laboratory and workflow specific criteria | Effective confirmation criteria can vary based on numerous details of a test’s methodology and its target genes. Changes can necessitate re-validation of confirmation criteria. |

**Abbreviations and definitions:**
SNV: Single Nucleotide Variant; Indel: Insertion or deletion; FP: False Positives; TP: True Positives; CI: 95% Confidence Interval; PPV: Positive Predictive Value; GIAB: Genome in a Bottle; GIAB-HC: Regions in which High-Confidence truth data are available for the GIAB specimens (unrelated to confidence in our own calls or to on/off-target regions).

Both laboratories’ bioinformatics pipelines have been described previously [8,19,20], although after these publications, both laboratories implemented the GATK Haplotype Caller [21,22] version 3.6 (Lab 1) or 3.7 (Lab 2). Lab 1 used a battery of criteria [8] to filter out clearly erroneous variants and to generate warnings on other variants that received manual review. Variants that failed review were also removed. To deliver high sensitivity, this process was conservative: Variants considered possibly true, despite warnings (e.g., relatively low depth or allele balance) were subjected to confirmation. Compared with Lab 1, Lab 2 used simpler filters (Quality-Depth (QD) score <4 and Fisher Strand bias (FS) score >40) and manual review was limited, resulting in a broader selection of variants being subjected to confirmation. Copy number and structural variants were excluded. As is typical for hybridization-based NGS, regions neighboring clinical targets received read coverage. In the GIAB specimens, variant calls within these “off-target” regions were used as long as they (a) passed the same quality filters as on-target calls, and (b) were within a set distance of a target (300 bp for Lab 1 and 50 bp for Lab parameters established in the clinical pipelines and not changed for this study). This requirement prevented large numbers of very low-coverage calls from being considered, although our quality filters remove most such calls in any case. In the clinical specimens, we were unable to use off-target calls because confirmatory data were unavailable.

Confirmation for clinical samples was performed by Lab 1 using Sanger (Thermo Fisher, Waltham, MA) or Pacific Biosciences (PacBio; Menlo Park, CA) amplicon sequencing. Lab 1 validated [23] the PacBio circular consensus sequencing method [24] specifically for use in confirmation. This method provides high accuracy [25] and has been successfully applied in other clinical genetic tests [26]. Lab 2 used only Sanger confirmation. When a confirmation assay and NGS disagreed, both results were manually reviewed, and if the reason for the disagreement was unclear, additional rounds of confirmation using different primers or assays were performed. There were few putatively mosaic variants in our study: Those present were considered TPs if confirmed and FPs if refuted by an adequately sensitive assay. Variants for which confirmation could not produce a confident answer (TP or FP) were not used in this study.

The GIAB reference calls [18] version 3.3.2 were used as truth data to confirm variants identified by each laboratory’s sequencing of the GIAB samples. Manual review was not performed for these samples. VCFeval [27] version 3.7.1 was used to compare each laboratory’s calls to the GIAB truth data to determine which laboratory calls were TPs and which were FPs. VCFeval can match variant calls even when the same diploid sequence is represented in different ways (“spelling differences”), an important factor for comparing indels and complex variants [27–29]. VCFeval also detects partial matches—i.e., zygosity errors (falsely calling a homozygous variant as heterozygous or vice versa) or heterozygous sites at which one of the called alleles is correct and one is not. These cases, reported by VCFeval as “FP_CA,” were considered FPs in our study (see Discussion). This study did not examine false negatives in detail.

Our analysis of the five GIAB specimens was restricted to the sample-specific high-confidence regions (GIAB-HC), which are annotated by the GIAB consortium to indicate where their reference data have high accuracy [18]. The GIAB-HC regions span 88-90% of the genome of each GIAB sample and cover most exons, introns, and intergenic regions, an improvement compared to older versions of the GIAB reference data [16] for which there was a greater bias toward “easy” regions [30]. The GIAB-HC designation is unrelated to any quality assessment of data produced by our laboratories. Indeed, our NGS assays produced both high- and low-quality variant calls within and outside the GIAB-HC regions. The GIAB-HC designation is also unrelated to whether calls were on- or off-target: most (not all) of our clinical targets were within the GIAB-HC regions, as were most off-target calls. Because the majority of our confirmatory data lay within the GIAB-HC regions, however, our confirmation criteria were selected by focusing within these regions (see Discussion). This required extrapolating the GIAB-HC regions to patient specimens, for which we used the union of the five GIAB-HC files—i.e., if a region was considered GIAB-HC in any of the five GIAB specimens, it was considered GIAB-HC in all patients. This approach prevented specific low-confidence calls in the reference data of particular GIAB specimen(s) from inappropriately annotating that site as low confidence in general.

Approximately two dozen quality metrics (Supplemental Table 1) were examined individually. The most useful metrics included different ways of measuring read depth (metrics A,B in the Supplemental Tables and Figures), allele balance for heterozygous (C) and homozygous calls (D), multiple quality scores (E,F), various indicators of strand bias (G-J), aspects of the variant call itself (K–N), and aspects of the genomic context (O). We manually chose candidate thresholds for each quantitative metric. Both the metrics and thresholds could be specific to a laboratory and variant type, but usually were not. Metrics for each variant call were then turned into discrete flags.

To delineate technically challenging genomic regions, we used the stratification BED (browser extensible data) files produced by the Global Alliance for Genomics and Health Benchmarking Workgroup [28]. These regions were padded by 10 bp on each side to ensure that all affected positions were appropriately annotated. These BED files were grouped as follows: (a) “repeats” combined homopolymers and short tandem repeats (STRs); (b) “segdups” (segmental duplications) included larger regions with homologous copies in the GRCh37 reference genome, and (c) “unmappable” regions were those in which our NGS reads could not map uniquely. Specific definitions are provided in Supplemental Table 1. The unmappable and segdup regions largely overlapped—but both were generally distinct from the (short) repeats.

These data were passed into a heuristic algorithm that selected a combination of flags for the final battery of criteria. The Python 2.7 code for this algorithm is available (LINK TO BE PROVIDED WITH FINAL DRAFT). Details of the algorithm are provided in the Supplemental Text.

Confidence intervals (CIs) were computed at 95% using the Jeffreys method. Both the Wilson score method and the tolerance interval method (as mentioned in the AMP/CAP guidelines) are included in Figure 2 for comparison.

**Figure 2:**
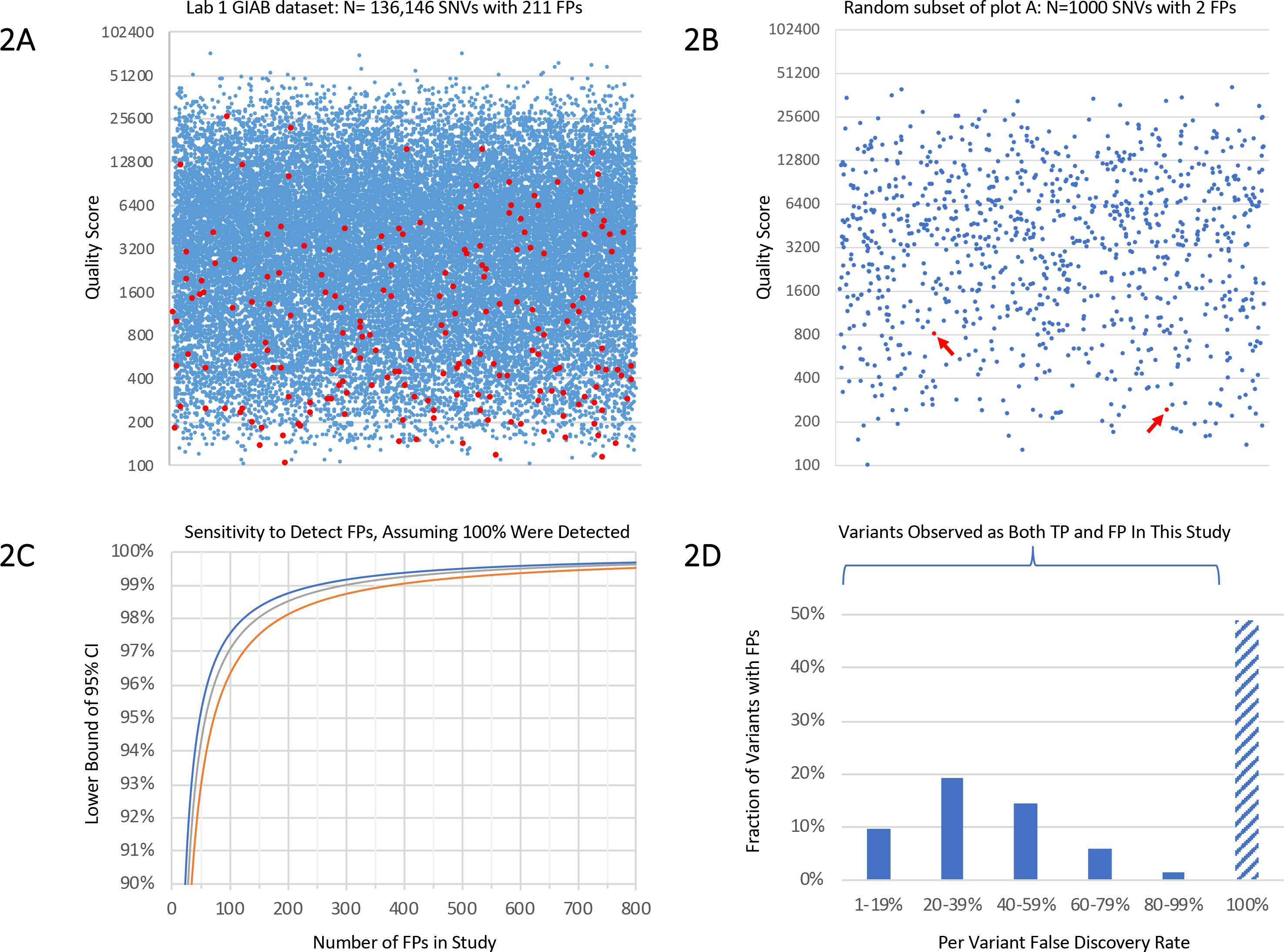
Individual quality metric results and statistics. (A) Variant call quality scores (QUAL, y-axis) for true positive (TP, blue) and false positive (FP, red) single-nucleotide variant (SNV) calls in the Lab 1 Genome in a Bottle (GIAB) dataset. The x-axis position of each point is randomly assigned. To make density changes visible, we plotted a random selection of one-fifth of the TP calls along with all FPs. In our large dataset, some FPs had quite high QUAL scores, demonstrating that this metric is inadequate alone. Corresponding histograms using the full dataset without down-sampling are shown in Supplemental Figure 1. (B) Random sample of 1000 data points from the same dataset as plot (A). All points are displayed. Arrows indicate the two FPs present. One thousand such random samples were generated, and compared with the full dataset in plot A, many would lead to quite different conclusions about the effectiveness of QUAL thresholds. (C) Lower bound of the 95% confidence interval (CI) on the fraction of FPs flagged (y-axis) as a function of the number of FPs used to determine criteria (x-axis) assuming 100% success is observed. The y-axis range is 90-100%. This calculation used the Jeffreys method (upper line, blue) the Wilson score method (bottom line, red), and the tolerance interval method (middle line, grey). All methods produce generally similar results and indicate the validity of any study such as ours. For example, using the Jeffreys method, flagging 49 of 49 FPs shows 100% effectiveness with a CI of 95% – 100%. Many prior studies did not achieve this level of statistical significance (Table 1). Consistent with these CI calculations, small datasets indeed resulted in ineffective criteria (see Results). (D) Histogram of per-variant false discovery rates (FDRs, x-axis) for all variants that (i) were observed more than once in our Lab 1 dataset, and (ii) for which one or more of those calls was an FP. SNVs and indels are combined. An FDR of 100% indicates a fully systematic FP (insofar as we can measure) and 0% a consistent TP (not shown in this graph). Each unique variant (i.e. a genetic alteration that may be present in multiple individuals) is counted once. The y-axis range is 0-50% of variants. Approximately half of all variants that were FPs were also correctly called as TPs in different specimen(s) or run(s). Examples of this were observed in both the clinical and GIAB specimens and included both SNVs and indels. Lab 2 results were similar. Many of these variants have low per-variant FDRs: These usually but not always are correctly called. Repeated TP observations of such a variant provide little information about the accuracy of any following observation of that same variant. Our study was underpowered to measure FDRs near 0% or 100%, and many more of these variants may exist than are shown here.

## Results

Our datasets included (a) almost 14,000 NGS variant calls subjected to confirmation during clinical testing, and (b) more than 184,000 calls with high-quality truth data from the GIAB samples (Table 1). We observed a total of 1662 FPs. To initially characterize our data. we calculated the false discovery rate (FDR) of calls in each set, defined as the number of FPs divided by the number of FPs plus TPs. Note that FDR is 1 minus the analytic positive predictive value (aPPV, also called positive percent agreement or PPA), a metric recommended by AMP/CAP guidelines, which consider it preferable to specificity (i.e., the specific calculation) for describing multi-gene sequencing tests [29]. As expected, the FDR of indels was considerably higher than that of SNVs (single nucleotide variants), and manual review had reduced FDRs in the Lab 1 clinical data compared to the GIAB data. FDRs in the GIAB samples were comparable between laboratories.

### Analysis of individual quality metrics from Lab 1

We examined a variety of metrics (Supplemental Tables 1 and 2, Supplemental Figures 1-4) to determine which were informative for identifying (or “flagging”) FPs. SNVs and indels were analyzed separately, as were the GIAB and clinical samples. For each metric and threshold, we computed the fraction of FPs and TPs flagged—these indicate the value and cost, respectively, of each potential flag. We also computed FDR, which indicates how likely flagged variants are to be FPs. We identified a number of informative metrics, although no single metric proved adequate alone. For example, consider the quality score (QUAL) for SNVs in the GIAB data (Figure 2A-B). This metric was used by Strom et al. [10] to identify SNVs that require confirmation, although Strom had only a single FP to use in defining thresholds (Supplemental Figure 5). In our, much larger dataset, no threshold value for QUAL could be chosen that flagged all, or even most, of the FPs without also flagging many TPs. Indeed, some FPs had quite high QUAL scores. A QUAL cutoff of 800, for example, flagged only 56% of FPs but also flagged 17% of TPs. These flagged variants had an FDR of 0.28%, indicating that this subset was not highly enriched for errors compared with the overall FDR of 0.15%. Indels fared similarly: 55% of FPs and 18% of TPs had QUAL<800, with an FDR of 16% compared with the baseline rate of 6.3%.

This observation was also true of our clinical dataset, in which 39% of FP SNVs and 2.9% of TPs had QUAL<800. These variants had a 3.6% FDR compared with 0.16% overall—20-fold enrichment—although most low-QUAL calls were still TPs. There is little precision in these measurements, however, because the clinical dataset contained only 10 FP SNVs, and only four had QUAL<800. Flagging all 10 FP SNVs required a QUAL threshold of 6400 (using logarithmic binning), which flagged almost 30% of TPs. Overfitting would certainly be an issue when analyzing the clinical dataset alone: If the single FP SNV with the highest QUAL score (5516) had been absent from these data, then thresholds would be driven by the next, much lower, score (2567). There is little statistical confidence in any such threshold: If 10 of 10 FPs are flagged, the point estimate is 100% FP sensitivity, but the lower bound of the CI is only 78% (Figure 2C), demonstrating that thresholds determined using such a small dataset could miss many FPs.

We similarly analyzed other metrics, finding that allele balance was the most informative, followed by strand bias, the presence of nearby variant calls, and certain characteristics of the variant itself (particularly, whether it was het-alt, meaning a heterozygous call in which neither allele is present in the GRCh37 reference genome). All of these criteria had limitations, however. Some provided strong indications of a call being an FP but flagged few such variants—strand bias was one example. More commonly, these criteria captured many TPs in addition to FPs yet still missed FPs at useful thresholds, similar to QUAL.

More than 80% of the Lab 1 GIAB variant calls were off-target. These calls passed our standard quality filters, although some had relatively low coverage (10–50x) or were in complex regions (repeats, high GC%, etc.). Such issues are present but less common within clinical targets. Nevertheless, the off-target data appeared representative: Indel FDRs were similar both on- and off-target (5.3% vs. 6.4%, respectively). SNV FDRs differed somewhat (0.04% vs. 0.17%) but remained low both on and off-target. Repeats accounted for half of the GIAB indels (both on- and off-target) and approximately 15% of the clinical indels. Repeats accounted for most of the indel FPs across our study although the vast majority of repeat-associated calls were correct, reflecting the high genetic variability of these sequences.

Of the variants observed more than once, approximately half of those with at least one FP call were also correctly called as TP in different runs(s) (Figure 2D). This was true for both SNVs and indels. It was even sometimes true across replicates of the same sample, a situation made possible because partial match errors were considered FPs in our analysis (see Methods). Many examples were found in the GIAB specimens, a result we attribute to the increased power we had to observe such variants in these data (see Supplemental Text). Clinical examples were observed as well. Case-by-case review suggests that the root causes of this behavior are varied and sometimes complex – partial match errors accounted for approximately half of these cases. Given our limited power to detect such variants, many more may exist. We conclude that historical confirmation performance is an ineffective quality metric in our datasets.

### Combining metrics

Because no single metric proved adequate, we investigated whether using multiple metrics might work better. One precedent for this approach is the Mu et al. study [12] which suggested that requiring depth>100 and allele balance between 40% and 60% would identify variants that do not require confirmation. In our dataset, which is much larger than Mu’s, this is not the case: 29 FP indels and seven FP SNVs failed to be flagged as requiring confirmation using Mu’s criteria. Nevertheless, we suspected that a larger battery of metrics might prove effective.

A heuristic algorithm was developed to explore this hypothesis. Briefly, this algorithm incrementally adds flags to a proposed set with the primary aim of capturing 100% of FPs using the combination of flags. This algorithm secondarily prioritizes minimizing the number of TPs also captured. Criteria were separately chosen for SNVs and indels. Variants in the GIAB and clinical samples were combined in this analysis to increase both the number and diversity of FPs available. This analysis was restricted to the GIAB-HC regions because of the limited amount of confirmatory data we had outside of these regions (see Discussion). To minimize overfitting, we performed Monte Carlo cross-validation, running the algorithm hundreds of times, in each iteration choosing flags (training) using two-thirds of the data (chosen at random) and testing these flags using the remaining one-third. We examined the flags selected by each iteration and the specific variants that led to differences among iterations (i.e., depending on whether the variant was randomly assigned to the training or the test set). This review highlighted the importance of particular implementation details of our algorithm (see Supplemental Text).

The final selected criteria are shown in Figure 3. Combined, these criteria flagged 100% of all 201 FP SNVs (CI 98.9–100%) and 100% of all 987 FP indels (CI 99.8–100%). Only 4.1% of clinical (i.e. non-GIAB) TP SNVs and 6.7% of TP indels were flagged by the same criteria. Many FPs received multiple flags, providing redundancy. Adding redundancy to our algorithm as an explicit objective increased the fraction of TPs flagged to 6.8% (SNVs) and 18% (indels). This increase was largely due to the addition of the repeat flag in this step (Supplemental Text).

**Figure 3:**
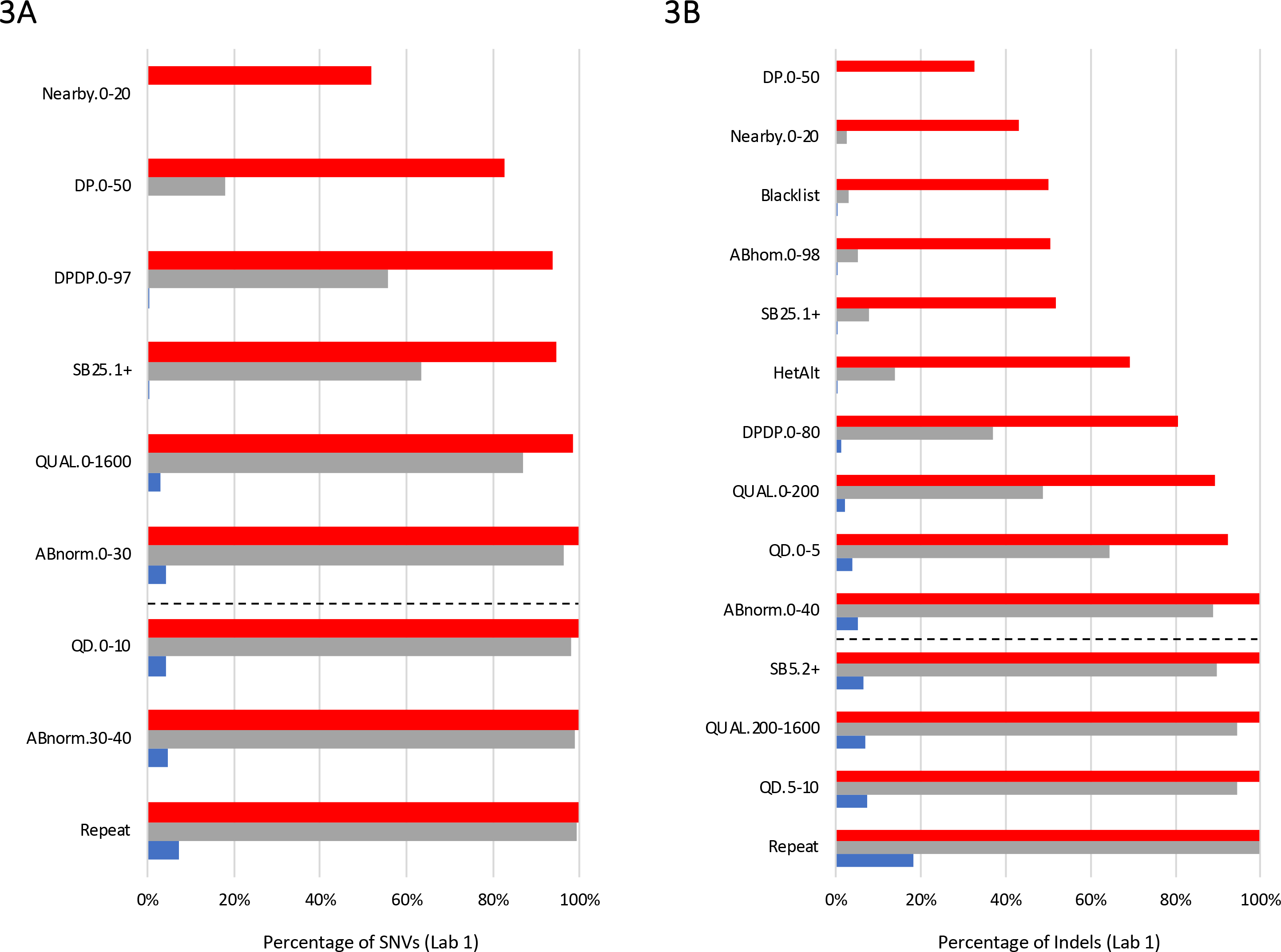
Combining flags. These plots show the cumulative effect (top to bottom) of sequentially combining flags chosen by our algorithm. Plot (A) shows single-nucleotide variants (SNVs), and plot (B) shows indels in the Lab 1 dataset. Red bars indicate the fraction of all false positives (FPs) captured and blue bars the fraction of clinical true positives (TPs). Grey bars show the fraction of FPs captured by two or more flags. The dotted lines illustrate the flags needed to capture 100% of the FPs using at least one flag each. To be conservative, we used the full set of flags shown here (maximizing double-coverage), and in particular requiring confirmation of all repeat-associated calls. Note that in the indel analysis, higher QUAL and QD thresholds were required to maximize double coverage than were required to achieve 100% capture of variants by a single flag each. The flags include:

DP: read depth, specifically GT_DP
DPDP: ratio of GT_DP to INFO_DP
SB5, SB25: strand bias metrics
QUAL: quality score
QD: quality-depth score
ABnorm: Allele balance for heterozygotes, normalized to be within 0.0-0.5
ABhom: Allele balance for homozygotes
HetAlt: Heterozygous call where neither allele is in the GRCh37 reference genome
Repeat: variant call within a homopolymer or short tandem repeat

We similarly assessed the importance of dataset size. For example, when we changed our crossvalidation to use only 50 FPs in training, then 100% of indel iterations and 71% of SNV iterations produced criteria that failed to flag at least some FPs in the test data. Between 2.0% – 6.0% of FP SNVs and 1.5% – 8.4% of FP indels were missed by the poorest performing 25% of datasets. These rates are consistent with our confidence interval calculations (Figure 2C), which predict that criteria established using such small datasets will capture between 95% – 100% of FPs, and will accomplish that 19 out of 20 times. Further reducing the number of training FPs to 20 produced criteria that uniformly failed, while 125 FP SNVs or 250 FP indels performed far better.

### Application of these methods to Lab 2

We examined whether a similar approach would work for the Lab 2 datasets, which were produced using somewhat different NGS methods. As of this analysis, Lab 2 had sequenced only one GIAB sample (NA12878) in addition to compiling clinical confirmation data. One consequence of this was that clinical FPs played larger role in determining both criteria and CIs. We do not consider this problematic, although we note that clinical confirmation data can have significant biases, such as over-representation of recurrent variants. Lab 2’s combined (clinical and GIAB) dataset was nevertheless diverse.

We first analyzed individual metrics (Supplemental Table 2), observing general similarities with Lab 1. Allele balance was the most informative criterion for separating TPs from FPs, with quality scores, read depth and strand bias also showing utility. Het-alt calls, variants with others nearby, and repeat-associated variants were often but not always FPs. As for Lab 1, none of these criteria was adequate alone. We combined flags (Supplemental Figure 6) and were able to capture 100% of FPs with CIs of 98.5–100% (SNVs) and 99.1–100% (indels). These criteria flagged 13.2% of TP SNVs and 15.4% of TP indels. requiring double-coverage increased these rates to 19.6% and 29.8%, respectively.

In comparison to Lab 1, the criteria chosen for Lab 2 appeared equally effective although less efficient—a greater fraction of TPs were flagged as requiring confirmation. One reason was that fewer “bad” variant calls had been removed prior to confirmation, a result of Lab 2’s different filtering and (for clinical specimens) manual review processes. This difference does not indicate an accuracy problem for Lab 2, but it did result in broader confirmation criteria. This observation reinforced our belief that confirmation criteria can vary based on filtering thresholds, and it supported our approach of first establishing (and validating) filters before establishing confirmation criteria. These results also suggest next steps for Lab 2: “Tightening up” filtering thresholds (where possible without impacting sensitivity) could further reduce confirmation workload by removing unambiguous FPs, similar to Lab 1. Reducing the number of FPs in this study, however, would make the CIs wider (i.e. less confident), reflecting an important design aspect of studies such as ours which depend on the set of FPs provided (see Discussion). The four additional GIAB samples would likely address this issue by providing additional, useful FPs.

## Discussion

Our study investigated whether a large and diverse dataset could be used to develop statistically robust criteria to guide the application of confirmation assays in clinical genetic testing. Our datasets combined clinical confirmation results with data from the sequencing of GIAB specimens, allowing for the two data types to complement each other, one key aspect of our methodology (Table 2). Similar to prior studies [10–14], our method identifies intermediate-confidence calls that require confirmation (i.e., to determine which are TPs and which are FPs) from calls that are high-confidence TPs from NGS alone and which do not benefit from confirmation. We cannot guarantee that the criteria chosen by our method (or any method) will capture all FPs. However, we can state that the probability of missing an FP is below a measurable level in datasets containing, collectively, almost 200,000 diverse variant calls with confirmatory data.

Our results differ from those of prior studies in important ways. Neither quality score (as suggested by Strom et al. [10]) nor the combination of allele balance and read depth (as suggested by Mu et al. [12]) captured all of the FPs in our datasets even when we re-optimized thresholds. Although we believe that criteria must be established independently for each NGS workflow—and indeed, workflows varied among these studies—we believe that this discrepancy results from the small datasets used in these prior studies, as well as their lack of separate training and test datasets. These limitations may have left these studies underpowered and subject to overfitting, which can impact effectiveness of the chosen criteria. A study by Baudhuin et al. [11] reported no FPs, a seemingly excellent result. However, our study and others [6,12,14] show that both FPs and TPs are abundant within the intermediate-confidence calls produced by current NGS methods. Aggressive filtering thus improves specificity at the expense of sensitivity. Indeed, a separate study showed that the methods used by Baudhuin had sensitivity limitations [7]. The two-threshold model (Figure 1A) helps address this issue.

In our datasets, a battery of criteria was required to flag FPs, consistent with metrics recommended by AMP/CAP guidelines [29]. This result is intuitive given that a variety of underlying factors can result in NGS FPs, and different FPs indeed have different properties. In theory, a single quality score that captures all of these factors could be simpler to use than a battery of criteria. Unfortunately, we know of no quality score produced by current variant callers that can identify FPs without also capturing many TPs. Other studies [14,21,31,32] agree.

A recent study by Van den Akker et al. [14] was somewhat similar to ours: These authors employed a supervised learning framework, used multiple quality metrics, and analyzed a larger dataset than was used in prior studies (albeit smaller than ours). There were important differences however: Van den Akker considered only one relatively small gene panel and omitted certain challenging regions of those genes. By contrast, we examined over 2000 genes and included off-target regions. Van den Akker’s logistic regression approach effectively creates (another) arbitrary quality score by mathematically combining metrics. We preferred to set individual thresholds on metrics recommended by guidelines [29], which makes our results understandable, easily implementable and avoids the statistical issues associated with mathematical combinations of highly correlated inputs.

We also preferred the objective of our heuristic algorithm, which focuses on the detection of FPs, as opposed to machine learning methods that optimize overall prediction accuracy, an approach that equally values the classification of TPs. Detecting FPs is clearly of paramount clinical importance. Moreover, in highly imbalanced datasets (i.e., every dataset in Table 1, in which TPs vastly outnumber FPs), such approaches can produce classifiers that work far better on the majority class (TPs) than on the minority (FPs) unless specific corrections are implemented [33]. Moreover, such approaches can be confounded by the many TPs with low quality metrics. Finally, overall performance metrics (e.g., prediction accuracy) for these methods also become uninformative with imbalanced data. For example, in our combined datasets (N = 194,119), an algorithm could show 99.6% prediction accuracy yet still miss half of the 1662 FPs. The FP-centric approach we used minimizes these issues.

Another important aspect of our methodology, not used in prior studies, was the inclusion of partial match errors, per recommendations [28]. These errors include both zygosity differences and incorrect diploid genotypes. Both of these error types can have significant clinical implications, and both are often resolved by confirmation. It is important to distinguish partial match errors from “spelling differences,” a different issue in which sequences are correct but are described in a non-canonical way [29]. Our study ignored spelling differences but considered partial match errors to be as important as “pure” FPs (i.e., cases in which a variant was called but none is actually present). Partial match errors represented approximately half of the FPs our GIAB datasets. Many partial match errors occurred in repeats, and thus they were less common in our clinical datasets (approximately 10% of FPs).

There were numerous benefits to using the large datasets we generated. These not only provided a diverse set of FPs for training (i.e., selecting criteria) but also allowed us to establish separate test sets to properly validate these criteria, identify outliers, and minimize overfitting. Using all five GIAB specimens also reduced any risk of biases resulting from the fact that variant callers are often trained, and may exhibit superior performance, on one of them (NA12878)[30]. Furthermore, our large datasets allowed us to compute statistical measures of confidence, a crucial part of any laboratory validation study, albeit one that is not always used [29]. The statistical metric we used (CI lower bound on fraction of FPs flagged) was inspired by recent guidelines [29,34]. It uses the size of the FP dataset as a proxy for whether those data are likely to be adequately representative and diverse for use in setting robust criteria. The CI bound does not directly measure diversity, however, and can be artificially inflated by under-filtering (see below). More sophisticated statistical approaches are ideal topics for future work. We caution against using metrics or statistics based on net accuracy of the classifier, which can be uninformative (as illustrated above), or PPV, which can be misleading because variants of intermediate quality become diluted by the large number of high-confidence TPs.

A key question is how large a dataset is adequate? For example, if certain criteria are shown to flag 49 of 49 FPs, these criteria are ostensibly 100% effective. However, they have only been statistically demonstrated to flag between 95% – 100% of FPs at p = 0.05 (i.e., 19 of 20 times) (Figure 2C). We demonstrated that this issue is not hypothetical: Small training datasets (e.g. 50 FPs) indeed resulted in poorly performing criteria, and we considered the corresponding CI bound (≥95.1%) inadequate. Our laboratories’ tests require high specificity, and we achieved 100% FP detection with CI lower bounds between 98.5% and 99.8% (Table 1) by using hundreds of example FPs. Such bounds are likely appropriate for panel and exome tests which have an increased risk of producing FPs compared to single-gene tests. Indels deserve careful attention, as FPs are more likely to appear pathogenic compared to FP SNVs.

Obtaining a large number of FPs for study can be challenging, and we were careful not to do so in artificial and problematic ways. For example, many FPs could be added by simply lowering filtering thresholds (under-filtering). However, the resulting criteria might be good only at flagging clearly erroneous calls as opposed to accurately defining the intermediate-confidence set (Figure 1A) for which confirmation matters most. This would be particularly problematic when using machine learning algorithms that best classify the largest input data subsets [33]. For example, Van den Akker’s FDRs are quite high [14] compared with those in our datasets and those of prior studies (Table 1) suggesting that many low-quality FPs may have been included. Under-filtering also will artificially tighten confidence intervals by counting obvious FPs, which could be misleading. A similarly problematic approach would be to run many samples containing the same FP variants. To avoid such problems, our study used the same filtering thresholds that our laboratories had previously validated for use in clinical practice, and we considered both the number of unique variants and the number of variant calls when designing our study (Table 1). The GIAB specimens provided a great deal of data, eliminating potential incentives to increase the dataset in problematic ways.

One might argue that our study artificially increased the number of FPs by including off-target regions. To the contrary, we considered it valuable to deliberately challenge our approach by adding off-target calls, some of which present technical challenges that are present, but less frequent in coding exons. Nevertheless, the off-target data appeared reasonably representative (see Results). Because we required 100% of the combined GIAB and clinical FPs to be flagged, we know that our criteria worked for both on- and off-target calls, and well as both GIAB and clinical specimens. We examined quality metrics separately for clinical, on-target GIAB, and off-target GIAB variant calls (Supplemental Table 2) observing similarities and expected differences resulting from (a) dataset sizes, (b) selection bias (in general, only clinically reportable variants were subject to confirmation in the clinical specimens), and (c) manual review (applied to the Lab 1 the clinical specimens but not GIAB).

Our study has important limitations. We used truth data provided by the GIAB consortium for confirmation in the GIAB specimens. These calls are highly accurate but imperfect [18]. In addition, when the Coriell GIAB specimens are sequenced (as we did), there can be genetic differences compared with the original DNA samples used to develop the truth data. In general, these issues will cause our methodology to establish broader criteria than might otherwise be necessary, increasing the number of TPs flagged. We examined outliers (FPs with unusually high confidence in our laboratory data) in the GIAB consortium’s data to ascertain whether those results appeared correct. We did not confirm any sites in the GIAB specimens using an orthogonal assay, although it may be valuable to do so in future studies.

A further limitation results from our focus on GIAB-HC regions, in which we had the largest numbers of variant calls with confirmatory data. The GIAB-HC designation does not indicate confidence in our own laboratories’ data (see Methods) although there is an important bias to recognize. The small fraction of the human genome (10-12%) that lies outside of the GIAB-HC regions are sites at which the GIAB consortium, using multiple platforms and extensive analysis, could not confidently determine the true sequence of their specimen(s) [16]. These are the “hardest” regions of the genome to sequence, and they present diverse challenges that increase error rates. Criteria that identify FPs within the GIAB-HC regions might not be effective outside of those regions, and we had little data to examine this. We concluded that clinically relevant variants outside of the GIAB-HC regions need to be confirmed, regardless of other quality metrics. A consequence of this was that variants in segmentally duplicated regions (which are usually not GIAB-HC) require confirmation. Within the Lab 1 dataset, 7.0% of clinically reported SNVs and 15% of clinical indels were thus flagged, in addition to those variants described in the Results above. Additional statistically valid studies would be required to determine which variant calls outside of the GIAB-HC regions could forgo confirmation in the future.

The ACMG guidelines for NGS recommend that laboratories have “extensive experience with NGS technology and be sufficiently aware of the pitfalls … before deciding that result confirmation with orthogonal technology can be eliminated” [3]. Our methodology provides a practical and rigorous way to follow this recommendation and to ensure that the “experience” (i.e., data) is based on a lab’s own specific methodologies and test targets, not NGS in general. Laboratories using the same sequencing instrument and variant caller (e.g. GATK) may still need different confirmation criteria owning to the many subtle differences between tests (particularly bioinformatics). Determining whether universal, interlaboratory criteria are possible would require additional, extensive study. Consistent with guidelines [3,5,29] our data suggest that each laboratory should validate its own confirmation criteria and that re-validation of these criteria should accompany any significant process change (including filtering changes).

Our results have specific implications for the current New York state guidelines, under which confirmation may be waived after “at least 10 positive samples per target gene” have been confirmed [4]. In our data, many FP variants were also called as TPs (Figure 2D), an issue not examined by prior studies. Examples were found in both the GIAB and clinical data from both laboratories. These variants run a high risk of being confirmed TPs in a series of tests and then, after confirmation is no longer considered necessary, being called falsely. Moreover, in our datasets, different variants within a gene often exhibited remarkably different properties that correlated with remarkably different FDRs. Observing some variants with a gene as TPs provides little information about whether other variants within that gene are FPs. In summary, our results argue against using the New York criteria for confirmation with multi-gene sequencing tests.

Our methodology does not address other roles that confirmation assays serve—e.g., verifying the identity of a specimen or determining the exact structure of certain variants. Furthermore, our framework does not directly address conflicts between NGS and confirmation assays. As Beck et al. [13] elegantly showed, naively assuming that a confirmation assay is always correct can introduce more errors than confirmation corrects. Our study also does not address the important issue of setting filtering criteria to ensure sensitivity. Laboratories need to address these issues separately. Finally, note that our approach is not necessarily optimal at minimizing the number of TPs that would receive confirmation. Instead, it is deliberately conservative and designed to avoid FPs escaping confirmation.

Our data show that criteria can be established to limit confirmation assays to a small fraction of variants without any measurable effect on analytic specificity. We show that a large and diverse dataset is required to accomplish this with confidence, and that the specifics of how criteria are chosen can have a substantial impact on their effectiveness (Table 2). Limiting confirmation assays in this careful manner may help reduce costs and improve the turnaround time of clinical genetic tests without compromising quality.

## Acknowledgements

We thank the Invitae and Partners HealthCare Laboratory for Molecular Medicine laboratory staff for producing the data used in this study. We thank all of the collaborators in the Genome in a Bottle Consortium for their excellent work on the reference specimens we used. We thank Nancy Jacoby for editorial help on this manuscript we thank Tina Hambuch and Katya Kosheleva for insightful comments.

## Notes

**Disclosures** Authors CL, ML, SL, RN, JP, HR, VR, and RT work for laboratories that provide clinical genetic testing services. SL, RN, JP, VR, and RT own stock and/or stock options in Invitae. SL owns stock in Illumina and Thermo Fisher. Certain commercial equipment, instruments, or materials are identified in this paper only to specify the experimental procedure adequately. Such identification is not intended to imply recommendation or endorsement by the National Institute of Standards and Technology, nor is it intended to imply that the materials or equipment identified are necessarily the best available for the purpose.

**Funding** All of the authors’ work on this project was funded by their respective employers. Grant funding was not used for this study.

## References

1. Korf BR, Rehm HL. New approaches to molecular diagnosis. JAMA, 2013, 309:1511–21

2. Rehm HL. Disease-targeted sequencing: a cornerstone in the clinic. Nat Rev Genet, 2013, 14:295–300

3. Rehm HL, Bale SJ, Bayrak-Toydemir P, Berg JS, Brown KK, Deignan JL, Friez MJ, Funke BH, Hegde MR, Lyon E, Working Group of the American College of Medical Genetics and Genomics Laboratory Quality Assurance Committee. ACMG clinical laboratory standards for next-generation sequencing. Genet Med, 2013, 15:733–47

4. New York State Department of Health. Guidelines for Validation Submissions of Next Generation Sequencing (NGS) assays under the NYS Testing Category of Genetic Testing, 2015

5. Aziz N, Zhao Q, Bry L, Driscoll DK, Funke B, Gibson JS, Grody WW, Hegde MR, Hoeltge GA, Leonard DGB, Merker JD, Nagarajan R, Palicki LA, Robetorye RS, Schrijver I, Weck KE, Voelkerding KV. College of American Pathologists’ laboratory standards for next-generation sequencing clinical tests. Arch Pathol Lab Med, 2015, 139:481–93

6. DePristo MA, Banks E, Poplin R, Garimella KV, Maguire JR, Hartl C, Philippakis AA, del Angel G, Rivas MA, Hanna M, McKenna A, Fennell TJ, Kernytsky AM, Sivachenko AY, Cibulskis K, Gabriel SB, Altshuler D, Daly MJ. A framework for variation discovery and genotyping using next-generation DNA sequencing data. Nat Genet, 2011, 43:491–8

7. Lincoln SE, Zook JM, Chowdhury S, Mahamdallie S, Fellowes A, Klee EW, Truty R, Huang C, Tomson FL, Cleveland MH, Vallone PM, Ding Y, Seal S, DeSilva W, Garlick RK, Salit M, Rahman N, Kingsmore SF, Aradhya S, Nussbaum RL, Ferber MJ, Shirts BH. An interlaboratory study of complex variant detection. bioRxiv, 2017:218529

8. Lincoln SE, Kobayashi Y, Anderson MJ, Yang S, Desmond AJ, Mills MA, Nilsen GB, Jacobs KB, Monzon FA, Kurian AW, Ford JM, Ellisen LW. A Systematic Comparison of Traditional and Multigene Panel Testing for Hereditary Breast and Ovarian Cancer Genes in More Than 1000 Patients. J Mol Diagn, 2015, 17:533–44

9. Mandelker D, Schmidt RJ, Ankala A, McDonald Gibson K, Bowser M, Sharma H, Duffy E, Hegde M, Santani A, Lebo M, Funke B. Navigating highly homologous genes in a molecular diagnostic setting: a resource for clinical next-generation sequencing. Genet Med, 2016, 18:1282–9

10. Strom SP, Lee H, Das K, Vilain E, Nelson SF, Grody WW, Deignan JL. Assessing the necessity of confirmatory testing for exome-sequencing results in a clinical molecular diagnostic laboratory. Genet Med, 2014, 16:510–5

11. Baudhuin LM, Lagerstedt SA, Klee EW, Fadra N, Oglesbee D, Ferber MJ. Confirming Variants in Next-Generation Sequencing Panel Testing by Sanger Sequencing. J Mol Diagn, 2015, 17:456–61

12. Mu W, Lu H-M, Chen J, Li S, Elliott AM. Sanger Confirmation Is Required to Achieve Optimal Sensitivity and Specificity in Next-Generation Sequencing Panel Testing. J Mol Diagn, 2016, 18:923–32

13. Beck TF, Mullikin JC, NISC Comparative Sequencing Program, Biesecker LG. Systematic Evaluation of Sanger Validation of Next-Generation Sequencing Variants. Clin Chem, 2016, 62:647–54

14. van den Akker J, Mishne G, Zimmer AD, Zhou AY. A machine learning model to determine the accuracy of variant calls in capture-based next generation sequencing. BMC Genomics, 2018, 19:263

15. Alpaydin E. Introduction to Machine Learning. MIT Press, 2014

16. Zook JM, Chapman B, Wang J, Mittelman D, Hofmann O, Hide W, Salit M. Integrating human sequence data sets provides a resource of benchmark SNP and indel genotype calls. Nat Biotechnol, 2014, 32:246–51

17. Zook JM, Catoe D, McDaniel J, Vang L, Spies N, Sidow A, Weng Z, Liu Y, Mason CE, Alexander N, Henaff E, McIntyre ABR, Chandramohan D, Chen F, Jaeger E, Moshrefi A, Pham K, Stedman W, Liang T, Saghbini M, Dzakula Z, Hastie A, Cao H, Deikus G, Schadt E, Sebra R, Bashir A, Truty RM, Chang CC, Gulbahce N, Zhao K, Ghosh S, Hyland F, Fu Y, Chaisson M, et al. Extensive sequencing of seven human genomes to characterize benchmark reference materials. Sci Data, 2016, 3:160025

18. Zook J, McDaniel J, Parikh H, Heaton H, Irvine SA, Trigg L, Truty R, McLean CY, De La Vega FM, Salit M, Genome in a Bottle Consortium. Reproducible integration of multiple sequencing datasets to form high-confidence SNP, indel, and reference calls for five human genome reference materials. bioRxiv, 2018:281006

19. Pugh TJ, Kelly MA, Gowrisankar S, Hynes E, Seidman MA, Baxter SM, Bowser M, Harrison B, Aaron D, Mahanta LM, Lakdawala NK, McDermott G, White ET, Rehm HL, Lebo M, Funke BH. The landscape of genetic variation in dilated cardiomyopathy as surveyed by clinical DNA sequencing. Genet Med, 2014, 16:601–8

20. Alfares AA, Kelly MA, McDermott G, Funke BH, Lebo MS, Baxter SB, Shen J, McLaughlin HM, Clark EH, Babb LJ, Cox SW, DePalma SR, Ho CY, Seidman JG, Seidman CE, Rehm HL. Results of clinical genetic testing of 2,912 probands with hypertrophic cardiomyopathy: expanded panels offer limited additional sensitivity. Genet Med, 2015, 17:880–8

21. Poplin R, Ruano-Rubio V, DePristo MA, Fennell TJ, Carneiro MO, Van der Auwera GA, Kling DE, Gauthier LD, Levy-Moonshine A, Roazen D, Shakir K, Thibault J, Chandran S, Whelan C, Lek M, Gabriel S, Daly MJ, Neale B, MacArthur DG, Banks E. Scaling accurate genetic variant discovery to tens of thousands of samples. bioRxiv, 2017:201178

22. McKenna A, Hanna M, Banks E, Sivachenko A, Cibulskis K, Kernytsky A, Garimella K, Altshuler D, Gabriel S, Daly M, DePristo MA. The Genome Analysis Toolkit: a MapReduce framework for analyzing next-generation DNA sequencing data. Genome Res, 2010, 20:1297–303

23. McCalmon S, Konvicka K, Reddy N, Olivares E, Whittaker J, Kautzer C, Rosendorff A. SMRTer Confirmation: Scalable clinical read-through variant confirmation using the Pacific Biosciences SMRT Sequencing Platform. American Society of Human Genetics 2016 Annual Meeting, Abstract 996F

24. Travers KJ, Chin C-S, Rank DR, Eid JS, Turner SW. A flexible and efficient template format for circular consensus sequencing and SNP detection. Nucleic Acids Res, 2010, 38:e159

25. Qiao W, Yang Y, Sebra R, Mendiratta G, Gaedigk A, Desnick RJ, Scott SA. Long-Read Single Molecule Real-Time Full Gene Sequencing of Cytochrome P450-2D6. Hum Mutat, 2016, 37:315–23

26. Ardui S, Race V, de Ravel T, Van Esch H, Devriendt K, Matthijs G, Vermeesch JR. Detecting AGG Interruptions in Females With a FMR1 Premutation by Long-Read Single-Molecule Sequencing: A 1 Year Clinical Experience. Front Genet, 2018, 9:150

27. Cleary JG, Braithwaite R, Gaastra K, Hilbush BS, Inglis S, Irvine SA, Jackson A, Littin R, Rathod M, Ware D, Zook JM, Trigg L, De La Vega FM. Comparing Variant Call Files for Performance Benchmarking of Next-Generation Sequencing Variant Calling Pipelines. bioRxiv, 2015:023754

28. Krusche P, Trigg L, Boutros PC, Mason CE, De La Vega FM, Moore BL, Gonzalez-Porta M, Eberle MA, Tezak Z, Labadibi S, Truty R, Asimenos G, Funke B, Fleharty M, Salit M, Zook JM, Global Alliance for Genomics and Health Benchmarking Team. Best Practices for Benchmarking Germline Small Variant Calls in Human Genomes. bioRxiv, 2018:270157

29. Roy S, Coldren C, Karunamurthy A, Kip NS, Klee EW, Lincoln SE, Leon A, Pullambhatla M, Temple-Smolkin RL, Voelkerding KV, Wang C, Carter AB. Standards and Guidelines for Validating Next-Generation Sequencing Bioinformatics Pipelines: A Joint Recommendation of the Association for Molecular Pathology and the College of American Pathologists. J Mol Diagn, 2018, 20:4–27

30. Li H, Bloom JM, Farjoun Y, Fleharty M, Gauthier LD, Neale B, MacArthur D. New synthetic-diploid benchmark for accurate variant calling evaluation. bioRxiv, 2017:223297

31. Pirooznia M, Kramer M, Parla J, Goes FS, Potash JB, McCombie WR, Zandi PP. Validation and assessment of variant calling pipelines for next-generation sequencing. Hum Genomics, 2014, 8:14

32. Goldfeder RL, Priest JR, Zook JM, Grove ME, Waggott D, Wheeler MT, Salit M, Ashley EA. Medical implications of technical accuracy in genome sequencing. Genome Med, 2016, 8:24

33. Batista GEAPA, Prati RC, Monard MC. A study of the behavior of several methods for balancing machine learning training data. ACM SIGKDD Explorations Newsletter, 2004, 6:20–9

34. Jennings LJ, Arcila ME, Corless C, Kamel-Reid S, Lubin IM, Pfeifer J, Temple-Smolkin RL, Voelkerding KV, Nikiforova MN. Guidelines for Validation of Next-Generation Sequencing-Based Oncology Panels: A Joint Consensus Recommendation of the Association for Molecular Pathology and College of American Pathologists. J Mol Diagn, 2017, 19:341–65

